# Taxonomic classification method for metagenomics based on core protein families with Core-Kaiju

**DOI:** 10.1101/2020.01.08.898395

**Authors:** Anna Tovo, Peter Menzel, Anders Krogh, Marco Cosentino Lagomarsino, Samir Suweis

## Abstract

Characterizing species diversity and composition of bacteria hosted by biota is revolutionizing our understanding of the role of symbiotic interactions in ecosystems. However, determining microbiomes diversity implies the classification of taxa composition within the sampled community, which is often done via the assignment of individual reads to taxa by comparison to reference databases. Although computational methods aimed at identifying the microbe(s) taxa are available, it is well known that inferences using different methods can vary widely depending on various biases. In this study, we first apply and compare different bioinformatics methods based on 16S ribosomal RNA gene and whole genome shotgun sequencing for taxonomic classification to three small mock communities of bacteria, of which the compositions are known. We show that none of these methods can infer both the true number of taxa and their abundances. We thus propose a novel approach, named Core-Kaiju, which combines the power of shotgun metagenomics data with a more focused marker gene classification method similar to 16S, but based on emergent statistics of core protein domain families. We thus test the proposed method on the three small mock communities and also on medium- and highly complex mock community datasets taken from the Critical Assessment of Metagenome Interpretation challenge. We show that Core-Kaiju reliably predicts both number of taxa and abundance of the analysed mock bacterial communities. Finally we apply our method on human gut samples, showing how Core-Kaiju may give more accurate ecological characterization and fresh view on real microbiomes.

## INTRODUCTION

Modern high-throughput genome sequencing techniques revolutionized ecological studies of microbial communities at an unprecedented range of taxa and scales (1, 2, 3, 4, 5). It is now possible to massively sequence genomic DNA directly from incredibly diverse environmental samples (3, 6) and gain novel insights about structure and metabolic functions of microbial communities.

One major biological question is the inference of the composition of a microbial community, that is, the relative abundances of the sampled organisms. In particular, the impact of microbial diversity and composition for the maintenance of human health is increasingly recognized (7, 8, 9, 10). Indeed, several studies suggest that the disruption of the normal microbial community structure, known as dysbiosis, is associated with diseases ranging from localized gastroenterologic disorders (11) to neurologic illnesses (12). However, it is impossible to define dysbiosis without first establishing what normal microbial community structure means within the healthy human microbiome. To this purpose, the Human Microbiome Project has analysed the largest cohort and set of distinct, clinically relevant body habitats (13), characterizing the ecology of healthy human-associated microbial communities. However there are several critical aspects. The study of the structure, function and diversity of the human microbiome has revealed that even healthy individuals differ remarkably in the contained species and their abundances. Much of this diversity remains unexplained, although diet, environment, host genetics and early microbial exposure have all been implicated. Characterizing a microbial community implies the classification of species/genera composition within the sampled community, which in turn requires the assignment of sequencing reads to taxa, usually by comparison to a reference database. Although computational methods aimed at identifying the microbe(s) taxa have an increasingly long history within bioinformatics (14, 15, 16), it is well known that inference based on 16S ribosomal RNA (rRNA) or shotgun sequencing vary widely (17). Moreover, even if data are obtained via the same experimental protocol, the usage of different computational methods or algorithm variants may lead to different results in the taxonomic classification. The two main experimental approaches for analyzing the microbiomes are based on 16S rRNA gene amplicon sequencing and whole genome shotgun sequencing (metagenomics).

Sequencing of amplicons from a region of the 16S rRNA gene is a common approach used to characterize microbiomes (18, 19) and many analysis tools are available (see Materials and Methods section). Besides the biases in the experimental protocol, a major issue with 16S amplicon-sequencing is the variance of copy numbers of the 16S genes between different taxa. Therefore, abundances inferred by read counts of the amplicons should be properly corrected by taking into account the copy number of the different genera detected in the sample (3, 20, 21). However, the average number of 16S rRNA copies is only known for a restricted selection of bacterial taxa. As a consequence, different algorithms have been proposed to infer from data the copy number of those taxa for which this information is not available (18, 22).

In contrast, whole genome shotgun sequencing of all the DNA present in a sample can inform about both diversity and abundance as well as metabolic functions of the species in the community (23). The accuracy of shotgun metagenomics species classification methods varies widely (24). In particular, these methods can typically result in a large number of false positive predictions, depending on the used sequence comparison algorithm and its parameters. For example in k-mer based methods as Kraken (25) and Kraken2 (26) the choice of *k* determines sensitivity and precision of the classification, such that sensitivity increases and precision decreases with increasing values for k, and vice versa. As we will show, false positive predictions often need to be corrected heuristically by removing all taxa with abundance below a given arbitrary threshold (see Materials and Methods section for an overview on different algorithms of taxonomy classification).

We highlight that the protocols for 16S-amplicons and shotgun methods are different and each has their own batch effects. Importantly, while shotgun taxonomic analysis gives classification results at species-level, 16S taxonomic profilers most often need to stop at the genus level. However, in the end, both aim at answering to the same question: “what are the relative abundances of taxa in the sample?” Therefore it is not methodologically wrong to compare their answers against the same community. To do that, it is possible to aggregate lower level (e.g. species) counts towards higher levels (e.g. genus), as it has been done in many benchmarks studies before (see, e.g., (17, 25, 27, 28)). In fact, several studies have performed comparisons of taxa inferred from 16S amplicon and shotgun sequencing data, with samples ranging from humans to studies of water and soil. Logares and collaborators (29) studied communities of bacteria marine plankton and found that shotgun approaches had an advantage over amplicons, as they rendered more truthful community richness and evenness estimates by avoiding PCR biases, and provided additional functional information. Chan et al. (30) analyzed thermophilic bacteria in hot spring water and found that amplicon and shotgun sequencing allowed for comparable phylum detection, but shotgun sequencing failed to detect three phyla. In another study (31) 16S rRNA and shotgun methods were compared in classifying community bacteria sampled from freshwater. Taxonomic composition of each 16S rRNA gene library was generally similar to its corresponding metagenome at the phylum level. At the genus level, however, there was a large amount of variation between the 16S rRNA sequences and the metagenomic contigs, which had a ten-fold resolution and sensitivity for genus diversity. More recently Jovel et al. (27) compared bacteria communities from different microbiomes (human, mice) and also from mock communities. They found that shotgun metagenomics offered a greater potential for identification of strains, which however still remained unsatisfactory. It also allowed increased taxonomic and functional resolution, as well as the discovery of new genomes and genes.

While shotgun metagenomics has certain advantages over amplicon-sequencing, its higher price point is still prohibitive for many applications. Therefore amplicon sequencing remains the go-to established cost-effective tool to the taxonomic composition of microbial communities. In fact, the usage of the 16S rRNA-gene as a universal marker throughout the entire bacterial kingdom made it easy to collect sequence information from a wide distribution of taxa, which is yet unmatched by whole genome databases. Several curated databases exist to date, with SILVA (32, 33), GreenGenes (34, 35) and Ribosomal Database Project (RDP) (36) being the most prominent. Additionally, NCBI also provides a curated collection of 16S reference sequences in its Targeted Loci project (https://www.ncbi.nlm.nih.gov/refseq/targetedloci/).

When benchmarking protocols for taxonomic classification from real samples of complex microbiomes, the “ground truth” of the contained taxa and their relative abundances is not known (see (27)). Therefore, the use of mock communities or simulated datasets remains as basis for a robust comparative evaluation of a method prediction accuracy. In the first part of this work we apply three widely used taxonomic classifiers for metagenomics, Kaiju (28), Kraken2 (26) and MetaPhlAn2 (37), and two common methods for analyzing 16S-amplicon sequencing data, DADA2 (38) and QIIME2 (39) to three small mock communities of bacteria, of which we know the exact composition (27). We show that 16S rRNA data efficiently allow to detect the number of taxa, but not their abundances, while shotgun metagenomics as Kaiju and Kraken2 give a reliable estimate of the most abundant genera, but the nature of the algorithms makes them predict a very large number of false-positive taxa.

The central contribution of this work is thus to develop a method to overcome the above limitations. In particular, we propose an updated version of Kaiju, which combines the power of shotgun metagenomics data with a more focused marker gene classification method, similar to 16S rRNA, but based on core protein domain families (40, 41, 42, 43) from the PFAM database (44).

Our criterion for choosing the set of marker domain families is that we uncover the existence of a set of core families that are typically at most present in one or very few copies per genome, but together cover uniquely all 8116 bacteria species in the PFAM database with an overall quite short sequence. Using presence of these core PFAMs (mostly related to ribosomal proteins) as a filter criterion allows for detecting the correct number of taxa in the sample. We tested our approach in a protocol called “Core-Kaiju” and show that it has a higher accuracy than other classification methods not only on the three small mock communities, but also on intermediate and highly biodiverse mock communities designed for the 1st Critical Assessment of Metagenome Interpretation (CAMI) challenge (45). In fact we will show how in all these cases Core-Kaiju overcomes, for the most part, the problem of false-positive genera and accurately predicts the abundances of the different detected taxa. We finally apply our novel pipeline to classify microbial genera in the human gut from the Human Macrobiome Project (HMP) (46) dataset, showing how Core-Kajiu may allow for a more accurate biodiversity characterization of real microbial communities, thus putting the basis for more solid dysbiosis analysis in microbiomes.

## MATERIALS AND METHODS

### Taxonomic classification: amplicon versus whole genome sequencing

Many computational tools are available for the analysis of both amplicon and shotgun sequencing data (25, 26, 28, 37, 38, 39, 47).

One of the differences among the several software for 16S rRNA analysis, is on how the next-generation sequencing error rate per nucleotide is taken into account, when associating each sampled 16s sequence read to taxa. Indeed, errors along the nucleotide sequence could lead to an inaccurate taxon identification and, consequently, to misleading diversity statistics.

The traditional approach to overcome this problem is to cluster amplicon sequences into the so-called operational taxonomic units (OTUs), which are based on an arbitrary shared similarity threshold usually set up equal to 97% for classification at the genus level. Of course, in this way, these approaches lead to a reduction of the phylogenetic resolution, since gene sequences below the fixed threshold cannot be distinguished one from the other.

That is why, sometimes, it may be preferable to work with exact amplicon sequence variants (ASVs), i.e. sequences recovered from a high-throughput marker gene analysis after the removal of spurious sequences generated during PCR amplification and/or sequencing techniques. The next step in these approaches is to compare the filtered sequences with reference libraries as those cited above. In this work, we chose to conduct the analyses with the following two opensource platforms: DADA2 (38) and QIIME2 (39). DADA2 is an R-package optimized to process large datasets (from 10s of millions to billions of reads) of amplicon sequencing data with the aim of inferring the ASVs from one or more samples. Once the spurious 16S rRNA gene sequences have been recovered, DADA2 allowed for the comparison with both SILVA, GreenGenes and RDP libraries. We performed the analyses for all the three possible choices. QIIME2 is another widely used bioinformatic platform for the exploration and analysis of microbial data which allows, for the sequence quality control step, to choose between different methods. For our comparisons, we performed this step by using Deblur (48), a novel sub-operational-taxonomic-unit approach which exploits information on error profiles to recover error-free 16S rRNA sequences from samples.

As shown in (27), where different amplicon sequencing methods are tested on both simulated and real data and the results are compared to those obtained with metagenomic pipelines, the whole genome approach resulted to outperform the previous ones in terms of both number of identified strains, taxonomic and functional resolution and reliability on estimates of microbial relative abundance distribution in samples.

Similar comparisons have also been performed with analogous results in (29, 30, 47, 49) (see (17) for a comprehensive summary of studies comparing different sequencing approaches and bioinformatic platforms).

Standard widespread taxonomic classification algorithms for metagenomics (e.g. Kraken (25) and Kraken2 (26)) extract all contained *k*–mers (all the possible strings of length *k* that are contained in the whole metagenome) from the sequencing reads and compare them with index of a genome database. However, the choice of the length *k* highly influences the classification, since, when *k* is too large, it is easy not to found a correspondence in reference database, whereas if *k* is too small, reads may be wrongly classified. Recently, a novel approach has been proposed for the classification of shotgun data based on sequence comparison to a reference database comprising protein sequences, which are much more conserved with respect to nucleotide sequences (28). Kaiju indexes the reference database using the Borrows-Wheeler-Transform (BWT), and translated sequencing reads are searched in the BWT using maximum exact matches, optionally allowing for a certain number of mismatches via a greedy heuristic approach. It has been shown (28) that Kaiju is able to classify more reads in real metagenomes than nucleotide-based *k*–mers methods. Therefore, previous studies on the community composition and structure of microbial communities in the human can be actually very biased by previous metagenomic analysis that were missing up to 90% of the reconstructed species (i.e. most of the species they found were not present in the gene catalog). We therefore chose to work with Kaiju (with MEM option (28)) for our taxonomic analysis. Although it resulted to give better estimates of sample biodiversity composition with respect to amplicon sequencing techniques, we found that it generally overestimates the number of genera actually present in our community (see Results section) of two magnitude orders, i.e. there is a long tail of low abundant false-positive taxa. To overcome this, we implemented a new release of the program, Core-Kaiju, which contains an additional preliminary step where reads sequences are firstly mapped against a newly protein reference library we created containing the amino-acid sequence of proteomes’ core PFAMs (see following section). We also compared standard Kaiju and Core-Kaiju results with those obtained via Kraken2 and via another widely used program for shotgun data analysis, MetaPhlAn2 (37, 47).

### Characterization of the core PFAM families

After downloading the PFAM database (version 32.0), we selected only bacterial proteomes and we tabulated the data into a *F* × *P* matrix, where each column represented a different proteome and each row a different protein domain. In particular, our database consisted of *P* = 8116 bacterial proteomes and *F* = 11286 protein families. In each matrix entry (*f,p*), we inserted the number of times the *f* family recurred in proteins of the *p* proteome, *n_f,p_*. By summing up over the *p* column, one can get the proteome length, i.e. the total number of families of which it is constituted, which we will denote with *l_p_*. Similarly, if we sum up over the *f* row, we get the family abundance, i.e. the number of times the *f* family appears in the PFAM database, which we call *a_f_*. Figure 1 shows the frequency histogram of the proteome sizes (left panel) and of the family abundances (right panel). Our primary goal was to find the so-called **core families** (50), i.e. the protein domains which are present in the overwhelming majority of the bacterium proteomes but occurring just few times in each of them (41, 51). In order to analyze the occurrences of PFAM in proteomes, we converted the original *F* × *P* matrix into a binary one, giving information on whether each PFAM was present or not in each proteome. In the left panel of Figure 2 we inserted the histogram of the family occurrences, which displays the typical *u-shape*, already observed in literature (43, 52, 53, 54): a huge number of families are present in only few proteomes (first pick in the histogram), whilst another smaller peak occurs at large values, meaning that there are also a percentage of domains occurring in almost all the proteomes. In the right panel, we show the plot of the number of rare PFAM (having abundance less or equal to four in each proteome) versus the percentage of proteomes in which they have been found. We thus selected the PFAMs found in more than 90% of the proteomes and such that max_*p*_*n*_*f,p*_ = 4 (see Zoom 2 panel of Figure 2).

**Figure 1.**
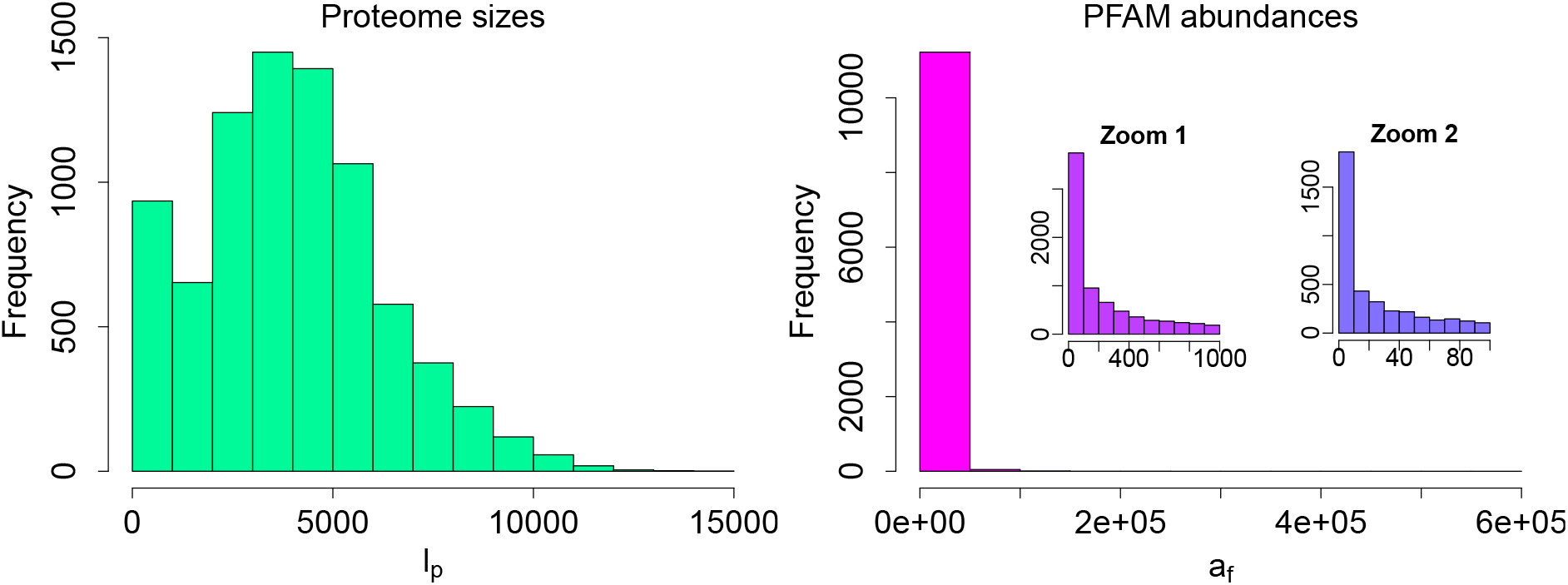
Proteome sizes and families abundances in PFAM database. On the left panel: frequency histogram of proteome lengths *l_p_* (total number of families of which a proteome *p* is composed). On the right panel: frequency histogram of family abundances, *a_f_* (number of times a PFAM *f* appears along a proteome).

**Figure 2.**
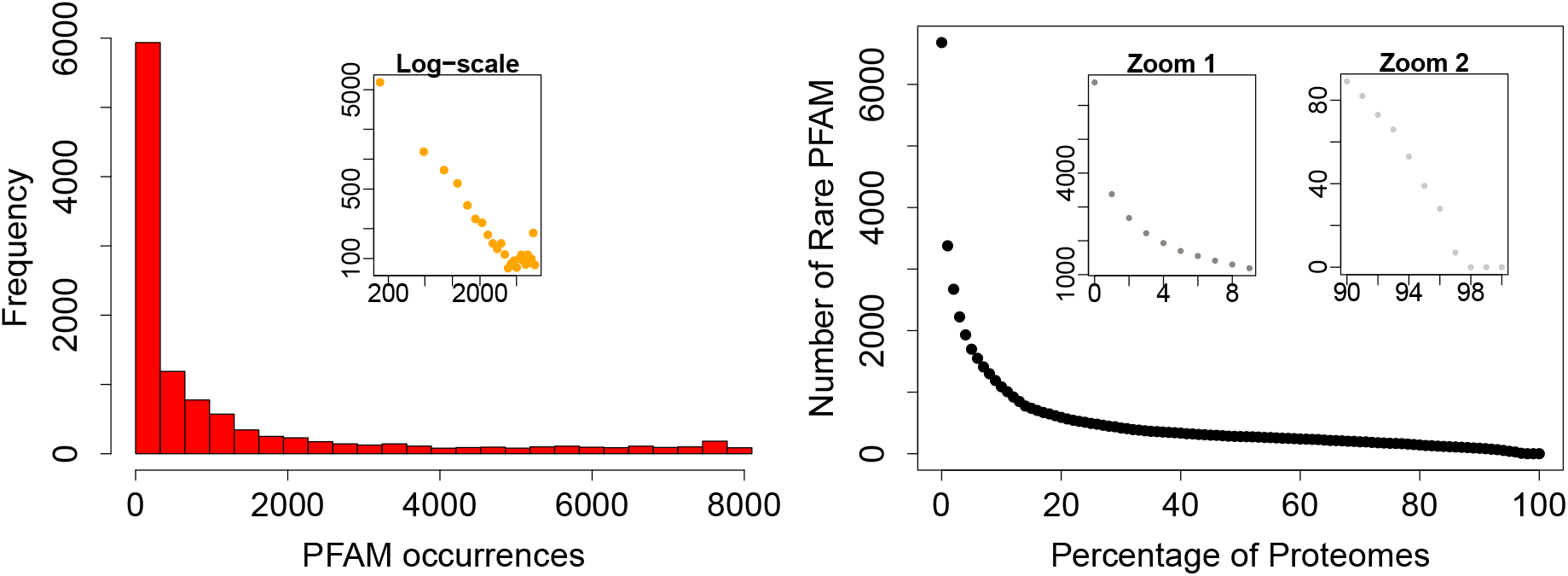
PFAM occurrences along proteomes. On the left panel: frequency histogram of family occurrences (number of proteomes in which a PFAM is contained). On the right panel: number of families with occurrence at most four versus the percentage of proteomes in which they are contained.

Since we wish to have at least one representative core PFAM for each proteome in the database, we checked whether with these selected core families we could ‘cover’ all bacteria. Unfortunately, none of them resulted to be present in proteomes 479430 and 1609106, corresponding to *Actinospica robiniae* DSM 44927 and *Streptomyces* sp. NRRL B-1568, respectively. We therefore looked for the most prevalent PFAM(s) present in such proteomes. We found that PFAM PF08338, occurring in 43% of the proteomes, was present in both *Actinospica robiniae* and *Streptomyces* and we therefore add it to our core-PFAM list. Eventually, in order to minimize the number of PFAMs to work with (and related computational cost), we considered in our final core-PFAM list only the minimum number of domains through which we were able to cover the whole list of proteomes of the databases. In particular, the selected core protein domains for bacteria proteomes are the ten PFAMs PF00453, PF00572, PF01029, PF01649, PF01795, PF03947, PF08338, PF09285 and PF17136 (see Table 1).

**Table 1.**
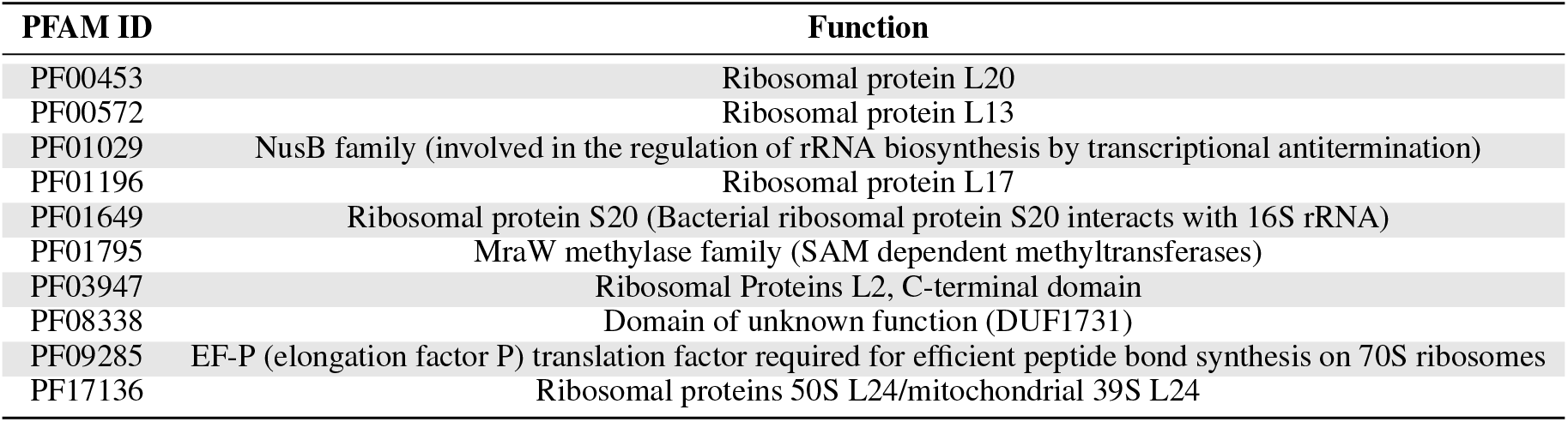
Core PFAMs identity number and corresponding function in proteomes.

#### Principal Coordinate Analysis

In order to explore whether the expression of the core PFAM protein domains are correlated with taxonomy, we did the following. First, we downloaded from the UniProt database (55) the amino acid sequence of each PFAM along the different proteomes (see Supporting Information for details). Their averaged (over proteomes) sequence lengths *L* resulted to be highly picked around specific values ranging from *L* = 46 to *L* = 297 (see Supporting Information, Figure S3, for the corresponding frequency histograms).

Second, for each family we computed the Damerau–Levenshtein (DL) distance between all its corresponding DNA sequences. DL measures the edit distance between two strings in terms of the minimum number of allowed operations needed to modify one string to match the other. Such operations include insertions, deletions/substitutions of single characters and transposition of two adjacent characters, which are common errors occurring during DNA polymerase. This analogy makes the DL distance a suitable metric for the variation between protein sequences. By simplicity and to have a more immediate insight, we conducted the analysis only for sequence points corresponding to the five most abundant phyla, i.e. Proteobacteria, Firmicutes, Actinobacteria, Bacteroidetes and Cyanobacteria.

After computing the DL distance matrices between all the amino-acid sequences of each PFAMs along proteomes, we performed the Multi Dimensional Scaling (MDS) or Principal Coordinate Analysis (PCoA) on the DL distance matrix. This step allow us to reduce the dimensionality of the space describing the distances between all pairs of core PFAMs of the different taxa and visualize it in a two dimensional space. In the last two columns of Table 2 we inserted the percentage of the variance explained by the first two principal coordinates for the ten different core families, where the first one ranges from 3.3 to 12.1% and the second one from 2.4 to 7.7%. We then plotted the sequence points into the new principal coordinate space, colouring them by phyla.

**Table 2.** Prevalence, Maximal/Total Occurences and Principal Coordinates of PFAM core families. We inserted, for each core family (PFAM ID, first column), the percentage of proteomes in which it appears (prevalence, second column), the maximum number of times it occurrs in one proteome (maximal occurrence, third column), the total number of times it is found among proteomes in the PFAM database (total occurrence, fourth column) and the percentage of variance explained by the firs two coordinates (PCo1 and PCo2, last two columns) when MDS is performed on sequences belonging to the five most abundant phyla (see Figure 3).

| PFAM ID | Prevalence | Maximal Occurrence | Total Occurrence | PCo1 | PCo2 |
| --- | --- | --- | --- | --- | --- |
| PF00453 | 95% | 3 | 7786 | 10.6% | 6.6% |
| PF00572 | 97% | 3 | 7897 | 5.4% | 5.1% |
| PF01029 | 96% | 4 | 12991 | 3.9% | 2.4% |
| PF01196 | 97% | 3 | 7888 | 12.1% | 5.7% |
| PF01649 | 94% | 3 | 7715 | 6.1% | 4.6% |
| PF01795 | 96% | 4 | 8113 | 5.2% | 4.9% |
| PF03947 | 97% | 4 | 7886 | 8.2% | 7.7% |
| PF08338 | 43% | 4 | 4267 | 3.3% | 2.9% |
| PF09285 | 96% | 4 | 8585 | 9.1% | 4.9% |
| PF17136 | 97% | 4 | 7896 | 5.4% | 4.1% |

In general, we observed a two-case scenario. For some families as PF03883 (see Figure 3, left panel), Actinobacteria and Proteobacteria sequences are grouped in one or two highly visible clusters each, whereas the other three phyla do not form well distinguished structures, being their sequence points close one another, especially for Cyanobacteria and Firmicutes. For other families as PF01196 (see Figure 3, left panel), all five phyla result to be clustered, suggesting a higher correlation between taxonomy and amino-acid sequences (see Supporting Information, Figure S4, for the other core families graphics). These results suggest that some core families (e.g. ribosomal ones) are phyla dependent, while other are not directly correlated with taxa.

**Figure 3.**
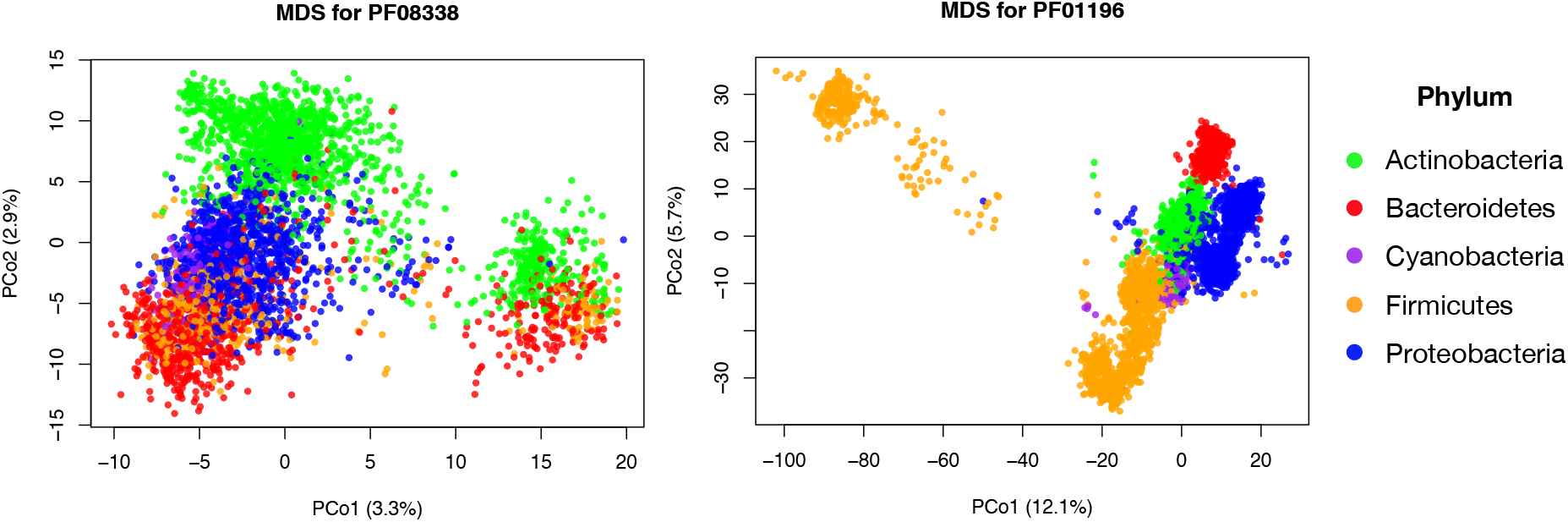
Phylum-based clustering for PF03883 and PF01196. For MDS analysis, only the sequences associated to the five most-abundant phyla (Proteobacteria, Firmicutes, Actinobacteria, Bacteroidetes, Cyanobacteria) have been considered.

### Mock bacteria communities

We started by testing shotgun versus 16S taxonomic pipelines on three small artificial bacterial communities generated by Jovel et al. (27), whose raw data are publicly available (Sequence Read Archive (SRA) portal of NCBI, accession number SRP059928). These mock populations contain DNA from eleven species belonging to seven genera: *Salmonella enterica, Streptococcus pyogenes, Escherichia coli, Lactobacillus helveticus, Lactobacillus delbrueckii, Lactobacillus plantarum, Clostridium sordelli, Bacteroides thetaiotaomicron, Bacteroides vulgatus, Bifidobacterium breve*, and *Bifidobacterium animalis*. For the taxonomic analysis at the genus level through 16S amplicon sequencing, we evaluated the performance of DADA2 (38) and QIIME2 pipelines (39). In particular, as shown in (27), QIIME2 produced more reliable results in terms of relative abundance of bacteria for all three mock communities when compared to Mothur (56), another widely used 16S pipeline, and to the MiSeq Reporter v2.5, a software developed by Illumina to analyze MiSeq instrument output data.

As for shotgun libraries, we tested the standard Kaiju (28), Kraken2 (26), the improved version of Kraken (25), and MetaPhlAn2 (37), the improved version of MetaPhlAn (47). This latter relies on unique clade-specific marker genes and it had been shown to have higher precision and speed over other programs (27).

Eventually, we tested Core-Kaiju on these mock communities and compared its performance with the above taxonomic classification methods.

### Core-Kaiju

After defining the core PFAMs, we created two protein databases for Kaiju: the first database only contains the protein sequences from the core families, whereas the second database is the standard Kaiju database based on the bacterial subset of the NCBI NR database. The protocol then follows these steps:

1. Classify the reads with Kaiju using the database with the core protein domains
2. Classify the reads with Kaiju using the NR database to get the preliminary relative abundances for each genus
3. Discard from the list of genera detected in (2) those having absolute abundance of less than or equal to twenty reads in the list obtained in point (1). This threshold represents our confidence level on the sequencing pipeline (see below).
4. Re-normalize the abundances of the genera obtained in point (3).

## RESULTS

### Comparison between methods, small mock community dataset

We evaluated the performance of both shotgun and 16S pipelines for the taxonomic classification of the three mock communities. In the top panels of Figure 4 we show the true relative genus abundance composition of the three small mock communities versus the ones predicted via the different tested taxonomic pipelines.

**Figure 4.**
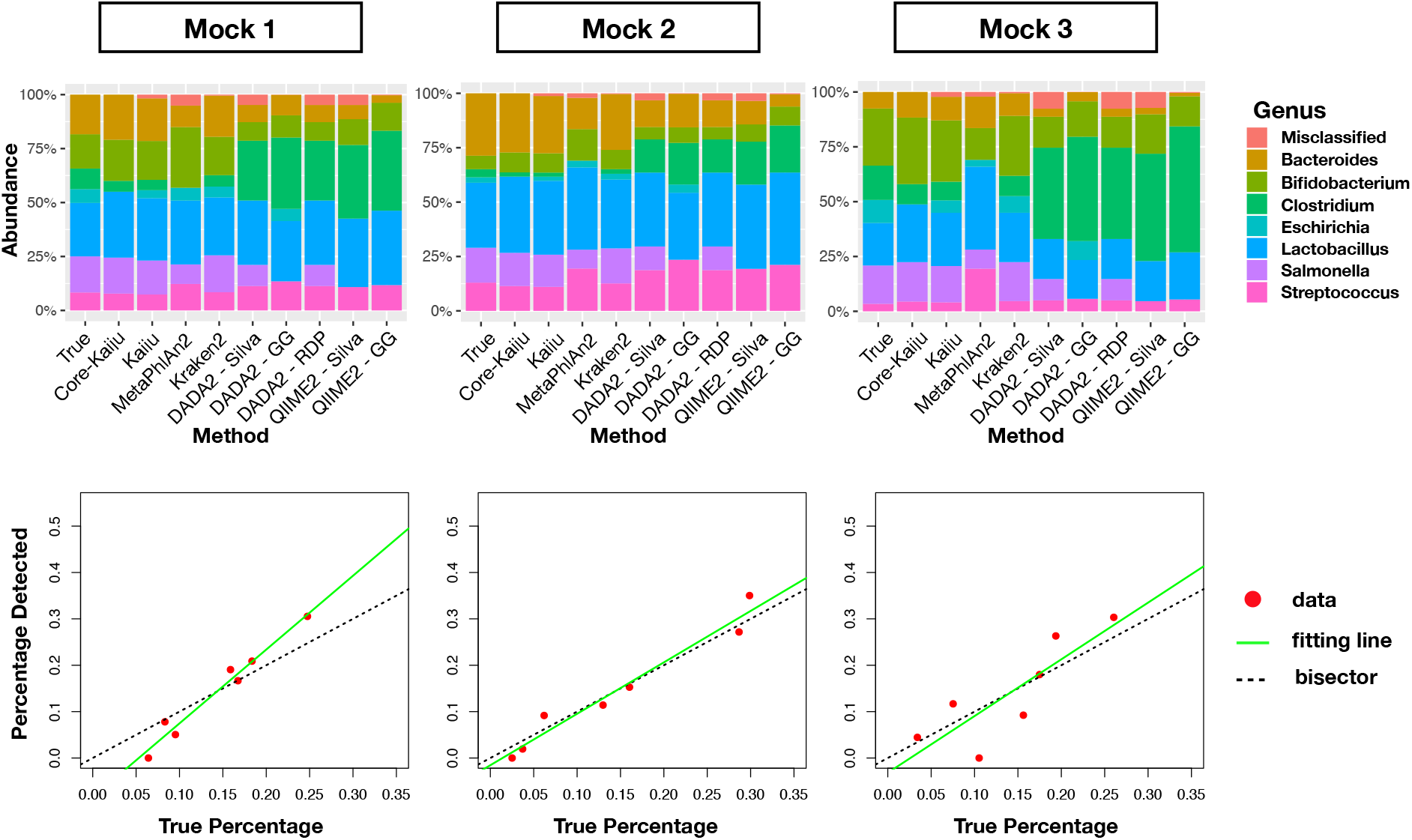
Comparison between theoretical and predicted relative abundances in small mock communities. **Top panels:** predicted relative abundance composition of the three small mock communities via different taxonomic classification methods. **Bottom panels:** red points represent data of relative abundance predicted for the genus level by Core-Kaiju on the three mock communities versus the true ones, known a priori. The green line is the linear fit performed on obtained points which, in the best scenario, should coincide with the quadrant bisector (dotted red line). In all three cases the predicted community composition was satisfactorily captured by our method, with an R-squared value of 0.97, 0.96 and 0.71, respectively.

We then applied the Core-Kaiju pipeline to detect the biodiversity composition of the same three mock communities. In Figure 4, bottom panels, we plot the linear fit performed on predicted relative abundances via Core-Kaiju versus theoretical ones, known a priori. As we can see, in all three cases the predicted community composition was satisfactorily captured by our method, with an *R*^2^ value higher than 0.7.

Our goal was to to quantitatively compare the performance of different methods in terms of both biodiversity and relative abundances. As for the first, we chose to measure it via the *F*_1_ score applied at the genera level. More precisely, we define the *recall* of a given taxonomic classification method as the number of truly-positive detected genera (present in a community and thus correctly detected by the method), *T_p_*, over the sum between *T_p_* and *F_n_*, the number of false-negative genera (present in a community, but missed to be classified). In contrast, we define the *precision* to be the ratio between *T_p_* and the sum of *T_n_* and *F_p_*, the number of false-positive genera (not present in a community and thus incorrectly detected as present). Finally, the *F*_1_ biodiversity score is twice the ratio between the product of recall and precision and their sum, i.e. 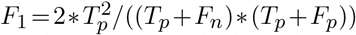. *F*_1_ score values obtained via the different methods for the three analysed mock communities are presented in Table 3. While *F*_1_ describes the overall accuracy in detecting the correct number of genera in the sample, *R*^2^ gives the correlation between the taxa abundance measured by the pipeline and the real composition of the microbial sample. Finally, we also indicated the number of genera each method predicts, *Ĝ*.

Table 3 summarizes the results of the analysis, together with the R-squared values, *R*^2^, obtained for the linear fit performed between true and predicted relative abundances. As we can see, both Core-Kaiju and MetaPhlAn2 gave a good estimate of the number of genera in the communities (which is equal to seven), whereas all 16S methods slightly overestimated it.

Finally, both standard Kaiju and Kraken2 predicted a number of genera much higher than the true one. Moreover, fit with standard Kaiju and Core-Kaiju of the predicted abundances displayed a higher determination coefficient with respect to all other pipelines, with the exception of Kraken2, which gave comparable values. However, if we focus on the *F*_1_ score, we can notice that Core-Kaiju outperformed all the other methods in terms of precision and recall. In particular, since the pipeline led to zero false-positive and only one false negative genus (*E.coli* in all three communities), the resulting precision and recall were 1 and 0.86 for all the sampled mocks. With Core-Kaiju, we were therefore able to produce a reliable estimate of both the number of genera within the communities and their relative abundances.

### Relative abundance vs absolute abundance thresholds

As stated in the introduction and observed above, metagenomic classification methods, such as Kaiju, often give a high number of false-positive predictions. In principle, one could set an arbitrary threshold on the detected relative abundances, for example 0.1% or 1%, to filter out low-abundance taxa that are likely false-positives. However, different choices of the threshold typically lead to very different results. The top panels of Figure 5 shows the empirical taxa abundance distribution of the 674 genera detected by Kaiju in the first small mock community. Such biodiversity number would decrease to 34, 9 or 7 if one considers only genera accounting for more than 0.01%, 0.1% and 1% of the total number of sample reads, respectively. Moreover, looking at the empirical pattern, one can notice the main gap between genera covering a fraction of less than 5 · 10^−3^ with respect to the total number of reads (black points) and those covering a fraction higher than 2 · 10^−2^ (green points), which corresponds to the genera actually present in the artificial community. One could therefore hope that, whenever such a gap is detected in the taxa abundance distribution, this corresponds to the one between false-positive and truly present taxa. However, as will be clear in the following section, this is not the case and it is not possible to set a relative threshold for the shotgun methods that works for all the mock communities.

**Figure 5.**
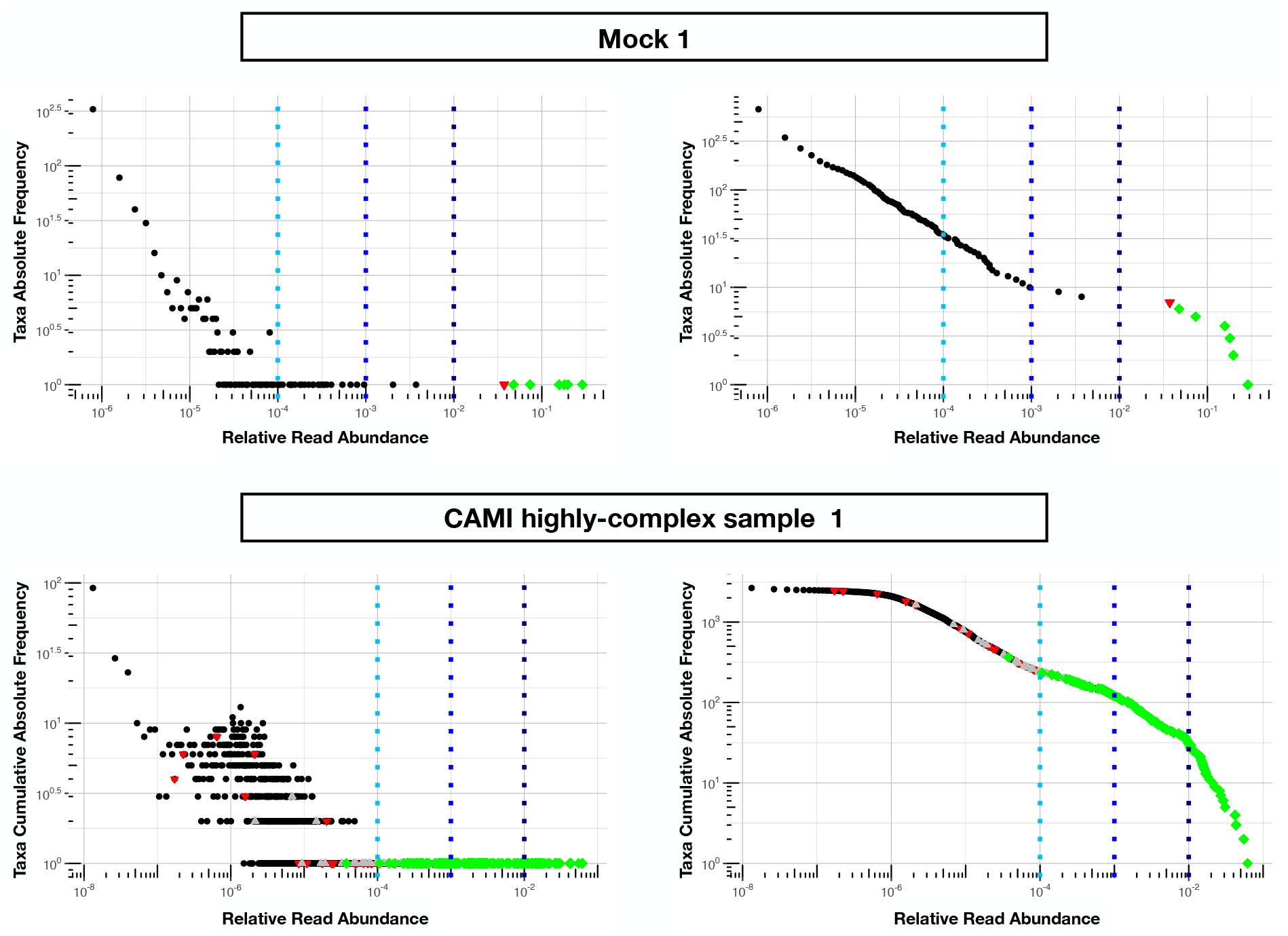
Relative vs absolute abundance thresholds for false-positive detection. **Top panels**: taxa abundance distribution plots for the first mock community (see Materials and Methods section). Green diamonds are the genera actually present in the artificial community and correctly detected by Core-Kaiju algorithm. The red triangle corresponds to the unique false-negative genus (*E.coli*) undetected with the newly proposed method. Dashed lines represent relative abundance thresholds on standard Kaiju output of 0.01%, 0.1% and 1%, respectively, which would have led to a biodiversity estimate of 34, 9 and 7 genera, respectively. Imposing an absolute abundance threshold of twenty reads on standard Kaiju output directly, would instead lead to an overestimation of 99 genera. **Bottom panels**: the same analyses have been performed on the CAMI high-complex sample 1. Again, green diamonds represent the 146 out of 193 genera present in the community and correctly detected by our pipeline. In this case, in addition to the remaining 47 false-negative genera (red triangles) we have also the presence of 58 false-negative genera, here represented by gray triangles. Setting a threshold on the relative abundance of reads produced by standard Kaiju gives a number of genera of 237 for the 0.01%, 120 for the 0.1% and 30 for the 1% threshold, respectively. Left and right panels represent, respectively, log-log absolute frequency and cumulative patterns of the taxa abundances in the mock communities.

**Table 3.** *F*_1_ score, R-squared values and number of predicted genera. For all three analysed mock communities, we inserted the *F*_1_ score (twice the ratio between the product of recall and precision and their sum), the *R*^2^ value of the linear fit performed between estimated and true abundances together with the number of predicted genera, *Ĝ*, with various taxonomic methods. The true number of genera is *G* = 7 for each community.

| | | Mock 1 ($G=7$) | | | Mock 2 ($G=7$) | | | Mock 3 ($G=7$) | | |
| --- | --- | --- | --- | --- | --- | --- | --- | --- | --- | --- |
| | | $F_1$ score | $R^2$ | $\hat{G}$ | $F_1$ score | $R^2$ | $\hat{G}$ | $F_1$ score | $R^2$ | $\hat{G}$ |
| Shotgun | Core-Kaiju | 0.92 | 0.97 | 6 | 0.92 | 0.96 | 6 | 0.92 | 0.71 | 6 |
|  | standard Kaiju | 0.02 | 0.97 | 674 | 0.03 | 0.98 | 501 | 0.02 | 0.94 | 738 |
|  | MetaPhlAn2 | 0.86 | 0.46 | 7 | 0.86 | 0.60 | 7 | 0.86 | 0.08 | 7 |
|  | Kraken2 | 0.04 | 0.98 | 333 | 0.05 | 0.99 | 266 | 0.04 | 0.96 | 378 |
| 16S | DADA2 + SILVA | 0.48 | 0.59 | 18 | 0.41 | 0.73 | 22 | 0.6 | 0.41 | 13 |
|  | DADA2 + GG | 0.5 | 0.45 | 17 | 0.43 | 0.60 | 21 | 0.63 | 0.35 | 12 |
|  | DADA2 + RDP | 0.48 | 0.59 | 18 | 0.4 | 0.73 | 23 | 0.6 | 0.41 | 13 |
|  | QIIME2 + SILVA | 0.21 | 0.50 | 0.21 | 41 | 0.59 | 41 | 0.21 | 0.43 | 41 |
|  | QIIME2 + GG | 0.26 | 0.46 | 32 | 0.26 | 0.50 | 32 | 0.25 | 0.36 | 33 |

### Application to CAMI challenge dataset

We tested and compared standard Kaiju, Kraken 2 and Core-Kaiju also on medium and high complexity mock bacterial communities obtained from the 1st CAMI challenge (45), in terms of biodiversity (recall, precision, *F*_1_ score, *Ĝ*) and abundance composition (linear fit R-squared). In Table 4 we show the results for samples 1 and 5 of the high-complexity dataset (see Supporting Information for the results of the other samples). As we can see, Core-Kaiju strongly outperformed the other methods in terms of precision. Indeed, it only slightly overestimated the true number of genera of around 10 taxa in sample 1, and 20 taxa in sample 5 (see Table 4), which is two order of magnitude lower with respect to the other methods (that predicted > 1600 of taxa). On the other hand, as also shown from the bottom panels of Figure 5, when using in standard Kaiju (or Kracken 2) a relative threshold of 1% so to reduce the number of false-positive taxa, as suggested by the previous analysis on the small mock community, the number of predicted taxa is in this case around 30, therefore strongly underestimating the real biodiversity of the samples.

As for the recall, the performance of Core-Kaiju (values around 77%) stands between standard Kaiju (values around 96%) and Kraken2 (values around 65%). The combination of recall and precision led to an *F*_1_ score around 74%, much higher than the other two pipelines (13%). Finally, as shown in Figure 6, Core-Kaiju gave also a very good estimation of the microbial composition, with an R-squared for the fit between theoretical and predicted relative abundances above 0.88, value comparable to standard Kaiju and much higher than the one obtained with Kraken2 (0.45). In the Supporting Information we present all the results for the other high-complexity samples as well as the analyses performed on the medium-complexity challenge dataset and the sensitivity of the classification on the absolute thresholds.

**Figure 6.**
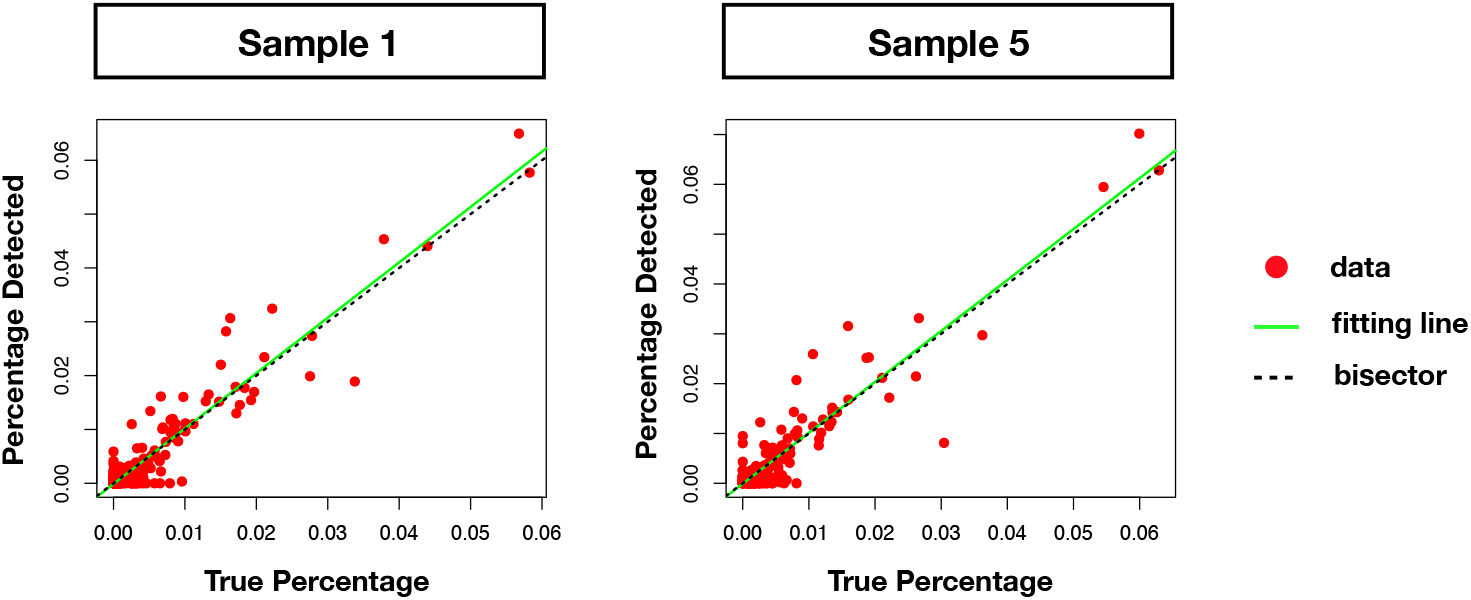
Linear fit between theoretical and predicted relative abundances with Core-Kaiju. red points represent data of relative abundance predicted for the genus level by Core-Kaiju on sample 1 and 5 from the CAMI highly-complex dataset versus the ground-truth abundances, known a priori. The green line is the linear fit performed on such values which, in the case of perfect matching between data and Cor-Kaiju output, should coincide with the quadrant bisector (dotted red line). In both cases, the predicted community composition was satisfactorily captured by our method, with a correlation with the real taxa abundances of *R*^2^ = 0.9 and *R*^2^ = 0.88 for sample 1 and 5, respectively.

### Application to human gut microbiome

We finally applied Core-Kaiju taxonomic classification method to an empirical data-set. We analysed a cohort of 26 healthy human fecal samples from the study (57) (metagenomic sequencing data are publicly available at the NCBISRA under accession number SRP057027). We applied standard Kaiju and found on average (over the 26 samples) 2108 bacterial genera. Similar overestimation of the number of taxa of Kajiu 1.0 would be obtained also with Kracken 2, highlighting the above mentioned problem of setting the correct threshold in order to have a realistic estimation of the sample biodiversity.

The right panel of Figure 7 shows the empirical taxa abundance distribution of one individual (sample ID: SRR2145359). As we can see, in this case the only apparent gap occurs between relative abundance of less than 10^−1^ and those above 0.5, with only one genus. It therefore results quite unrealistic that all the taxa but one should be considered falsepositive. The same plot shows the vertical lines corresponding to threshold on relative population of 0.01%, 0.1% and 1% above which we have 97, 32 and 10 taxa, respectively.

**Figure 7.**
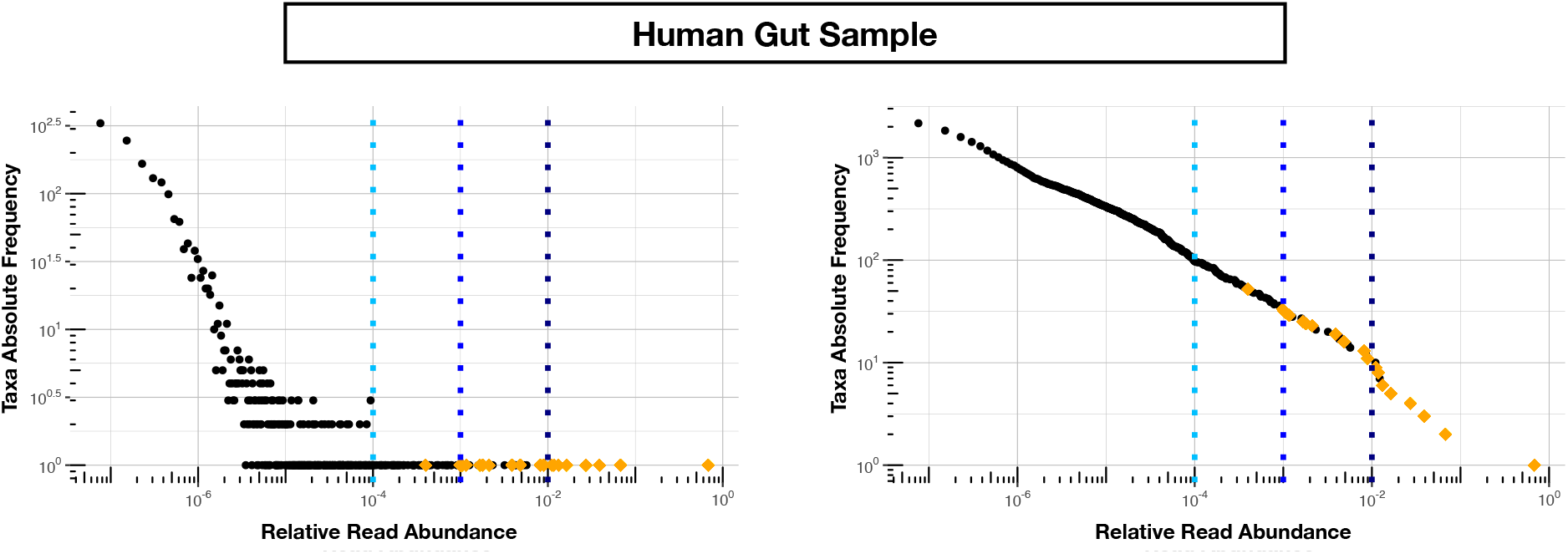
Relative vs absolute abundance thresholds in the human gut sample. Taxa abundance distribution plots for a human gut sample of a healthy individual, where standard Kaiju detects (without any threshold) 2165 genera. In this case the number (and label) of the actual present genera is unknown. Nevertheless estimates from a reference cohort of stool microbiomes (46) from 174 healthy HMP participants (16S V3-V5 region, >5k reads per sample, 97% OTU clustering), report an average number of genera per sample of 25 (max=46, min=9) (1). Setting a threshold on the relative abundance of reads produced by standard Kaiju gives a number of genera of 97 for the 0.01%, 32 for the 0.1% and 10 for the 1% threshold, respectively. In contrast, considering false-positive all genera with less or equal to twenty reads in standard Kaiju output, we end up with 625 genera. Orange diamonds in plot correspond to the 21 genera detected with Core-Kaiju, a number compatible with the reported estimates. Left and right panels represent log-log absolute frequency and cumulative patterns, respectively.

In contrast, with Core-Kaiju we did not need to tune a relative threshold. Instead, by removing false-positive through the (fixed) absolute abundance of 20 reads we ended up with 21 genera (orange diamonds in Figure 7), which is compatible with previous estimates. In fact, the available ampliconsequencing datasets from stool samples of healthy participants of the human microbiome project (1) suggest that there are on average 25 different bacterial genera per sample (based on 174 samples with at least >5k reads per sample using 97% OTU clustering). However, in terms of taxa composition, Core-Kaiju predicted abundances are different from those obtained using 16s classification methods (1).

## DISCUSSION

An important source of errors in the performance of any algorithm working on shotgun data is the high level of plasticity of bacterial genomes, due to widespread horizontal transfer (41, 58, 59, 60, 61, 62). Indeed, most highly abundant gene families are shared and exchanged across genera, making them both a confounding factor and a computational burden for algorithms attempting to extract species presence and abundance information. Thus, while having access to the sequences from the whole metagenome is very useful for functional characterization, restriction to a smaller set of families may be a very good idea when the goal is to identify the species taxa and their abundance.

To summarize, we have presented a novel method for the taxonomic classification of microbial communities which exploits the peculiar advantages of both whole-genome and 16S rRNA pipelines. Indeed, while the first approaches are recognised to better estimate the relative taxa composition of samples, the second are much more reliable in predicting the true biodiversity of a community, since the comparison between taxa-specific hyper-variable regions of bacterial 16S ribosomal gene and comprehensive reference databases allows in general to avoid the phenomenon of false-positive taxa detection. Indeed, the identification of a threshold in shotgun methods to remove most of the false-positive is of course a critical problem, because in general the true taxa composition is not known, and thus setting the wrong threshold may lead to a huge over- (or under-) estimation of the sample biodiversity, as shown in this work.

**Table 4.** Performance comparison on CAMI high-complexity samples 1 and 5. In the first four columns, we inserted the values for the precision, the recall, the *F*_1_ score, the *R*^2^ value of the linear fit performed between estimated and true abundances, and the number of predicted genera *Ĝ* with Core-Kaiju, standard Kaiju and Kraken2. The true number of genera is *G* = 193 for each sample. In the last column we also inserted the number of genera one would predict with standard Kaiju and Kraken2 by setting a relative threshold of 1%, i.e. by considering false-positive all those genera having a relative abundance of less than 0.01 in the sample. We denoted this quantity by *Ĝ*_1%_.

| | Sample 1 ($G = 193$) | | | | | | Sample 5 ($G = 193$) | | | | | |
| --- | --- | --- | --- | --- | --- | --- | --- | --- | --- | --- | --- | --- |
| | Precision | Recall | $F_1$ score | $R^2$ | $\hat{G}$ | $\hat{G}_{1\%}$ | Precision | Recall | $F_1$ score | $R^2$ | $\hat{G}$ | $\hat{G}_{1\%}$ |
| **Core-Kaiju** | 0.72 | 0.76 | 0.74 | 0.90 | 204 | — | 0.72 | 0.79 | 0.75 | 0.88 | 213 | — |
| **standard Kaiju** | 0.07 | 0.96 | 0.13 | 0.92 | 2652 | 30 | 0.07 | 0.96 | 0.13 | 0.89 | 2660 | 26 |
| **Kraken2** | 0.07 | 0.65 | 0.13 | 0.45 | 1715 | 27 | 0.07 | 0.65 | 0.13 | 0.45 | 1697 | 26 |

Inspired by the role of 16S gene as a taxonomic fingerprint and by the knowledge that proteins are more conserved than DNA sequences, we proposed an updated version of Kaiju, an open-source program for the taxonomic classification of whole-genome high-throughput sequencing reads where sample metagenomic DNA sequences are firstly converted into amino-acid sequences and then compared to microbial protein reference databases. We identified a class of ten domains, here denoted by core PFAMs, which, analogously to 16S rRNA gene, on one hand are present in the overwhelming majority of proteomes, therefore covering the whole domain of known bacteria, and which on the other hand occur just few times in each of them, thus allowing for the creation of a novel reference database where a fast research can be performed between sample reads and PFAMs amino-acid sequences. Tested against mock microbial communities, of different level of complexity, generated in other studies (27, 45) and available online, the proposed updated version of Kaiju, Core-Kaiju, outperformed popular 16S rRNA and shotgun methods for taxonomic classification in the estimation of both the total biodiversity and taxa relative abundance distribution. In fact, by fixing an absolute threshold with Core-Kaiju (by only considering abundances greater to twenty reads), we are able to correctly classify the biodiversity in all samples of different size and complexity, while keeping a very good performance in the prediction of taxa abundances.

We highlight that other technologies exist beyond metagenomics or 16S amplicons on a MiSeq (integrated instrument performing clonal amplification and sequencing), as for example PaCBio (63). Earl and collaborators (64) used a CAMI dataset to test the accuracy of this method and it is therefore possible to indirectly compare Core-Kaiju with PaCBio through their results. Also in this case we found that our method gives a slightly higher *R*^2^ score for the genera abundances composition, confirming the competitiveness of Core-Kaiju even with long-read technology such as PaCBio. However, a deeper comparison with these methods goes beyond the scope this work because, although might perform better than MiSeq next-generation sequencing approaches, they are quite rare and available only for much higher price.

Our promising results pave the way for the application of the newly proposed pipeline in the field of microbiota-host interactions, a rich and open research field which has recently attracted the attention of the scientific world due to the hypothesised connection between human microbiome and healthy/disease (65, 66). Having a trustable tool for the detection of microbial biodiversity, as measured by the number of genera and their abundances, could have a fundamental impact in our knowledge of human microbial communities and could therefore lay the foundations for the identification of the main ecological properties modulating the healthy or ill status of an individual, which, in turn, could be of great help in preventing and treating diseases on the basis of the observed patterns.

## DATA AVAILABILITY

All data and codes used for this study are available online or upon request to the authors. Raw data for the three in-silico mock communities (27) are publicly available at the Sequence Read Archive (SRA) portal of NCBI under accession number SRP059928. Metagenomic sequencing data of the healthy human fecal samples from the study (57) are publicly available at the NCBI SRA under accession number SRP057027. CAMI medium and high complexity datasets are available at https://data.cami-challenge.org/participate under request.

## FUNDING AND ACKNOWLEDGEMENTS

This work was supported by the STARS GRANT UNIPD ReACT to S.S. MCL, S.S. and A.K. acknowledge Cariparo foundation Visiting Program 2018.

